# Constrained Laser-Induced Cavitation for Miniaturized Treatment of Deep Vein Thrombosis

**DOI:** 10.64898/2026.08.14.744960

**Authors:** Joseph Yang, Daiwei Li, Kevin Wang, Pei Zhong, Junjie Yao

## Abstract

Chronic, mechanically resilient thrombi remain difficult to remove rapidly and safely using existing therapies, which are limited by slow treatment speeds, reduced efficacy against aged clots and risks associated with embolic debris. Here we introduce Constrained Laser-Induced Cavitation (CLIC), a novel approach that confines laser-induced cavitation bubble generation and collapse within a miniaturized waveguide to enhance thrombolysis. Optimized CLIC removed retracted clots at a mass-loss rate of 393.5 mg/min, ∼40-fold higher than reported state-of-the-art sonothrombolysis under similar conditions. Systematic variation of channel length and laser parameters showed that CLIC efficacy depends strongly on treatment geometry and cavitation dynamics. Post-treatment analysis revealed cylindrical channels consistent with clot removal dominated by fluid jetting and suction-driven evacuation, with cavitation shockwaves likely contributing a secondary role. Debris fragment measurements remained predominantly below a 1 mm embolic-risk threshold, consistent with a promising embolic safety profile. These findings establish CLIC as a viable strategy for rapid thrombolysis of chronic, mechanically resistant thrombi.

## 1. Introduction

Deep vein thrombosis (DVT) describes the formation of thrombi, also known as blood clots, in deep venous circulation. DVT is a high-morbidity, high-occurrence pathology that is the third-highest cause of cardiovascular death/disability in the US [1, 2], affecting 2.5 million individuals annually in the US alone [3], with 1 in 3 patients developing a recurrent DVT within 5 years [4]. A typical subacute venous thrombus consists of tightly packed red blood cells within a fibrin network, alongside some platelets and leukocytes [5–7]. Treatment of thrombi typically aims to disrupt this fibrin structure while minimizing the size of released thrombus debris to avoid distal emboli [8–11]. For treatment of high-risk, typically aged thrombi, the American Heart Association recommends pharmacological dissolution (use of thrombolytic agents such as tPA) or mechanical fragmentation (i.e., physical removal of the clot). However, these contemporary DVT treatments are limited by low efficacy (∼20% failure rate), long procedure times (hours to days), and major intraoperative risks, e.g., major bleeding/intracranial bleeding (9.2% and 1.46% respectively) [12, 13] and distal emboli [3, 14].

A largely unexplored, photonics-based approach to thrombolysis is short-pulse laser-induced cavitation, best known from laser lithotripsy for kidney stones [15, 16]. Laser-induced cavitation typically uses long-wavelength light (e.g., 2080 nm) matched to the absorption peaks of water. A high-energy laser pulse, typically on the levels of 0.2 to 1 Joules per pulse (Jpp), is used to vaporize the fluid in front of an optical fiber. The vaporized water generates an unconstrained vapor bubble that collapses into a toroidal bubble, inducing a shockwave [16–18]. Current understanding attributes kidney-stone treatment primarily to the shockwave generated by vapor-bubble collapse [15, 16, 18] rather than thermal ablation. Based on this understanding, we applied laser-induced cavitation to thrombi, adding a waveguide with the initial goal of redirecting shockwave energy towards the target and protecting surrounding vasculature from off-target shockwaves. One additional advantage of an optical fiber-based approach is that it enables substantially smaller catheters than conventional thrombolysis modalities such as sonothrombolysis [19, 20] while still allowing for important surgical capabilities, such as the aspiration of debris or injection of exogenous agents for treatment efficacy or imaging guidance [21, 22]. Smaller catheters allow treatment of thrombi in narrower vessels and may improve post-operative outcomes by reducing risks such as vascular damage during catheter insertion [23, 24].

The conceptual framework of CLIC is illustrated in Fig. 1. An optical fiber is positioned within a catheter equipped with a stainless-steel waveguide and advanced to the target thrombus. Adding the waveguide substantially altered both treatment efficacy and the underlying mechanisms of CLIC. The effective cavity length (ECL), which describes the distance between the optical fiber tip and waveguide tip, must be optimized during treatment to ensure most efficient cavitation dynamics.

**Figure 1:**
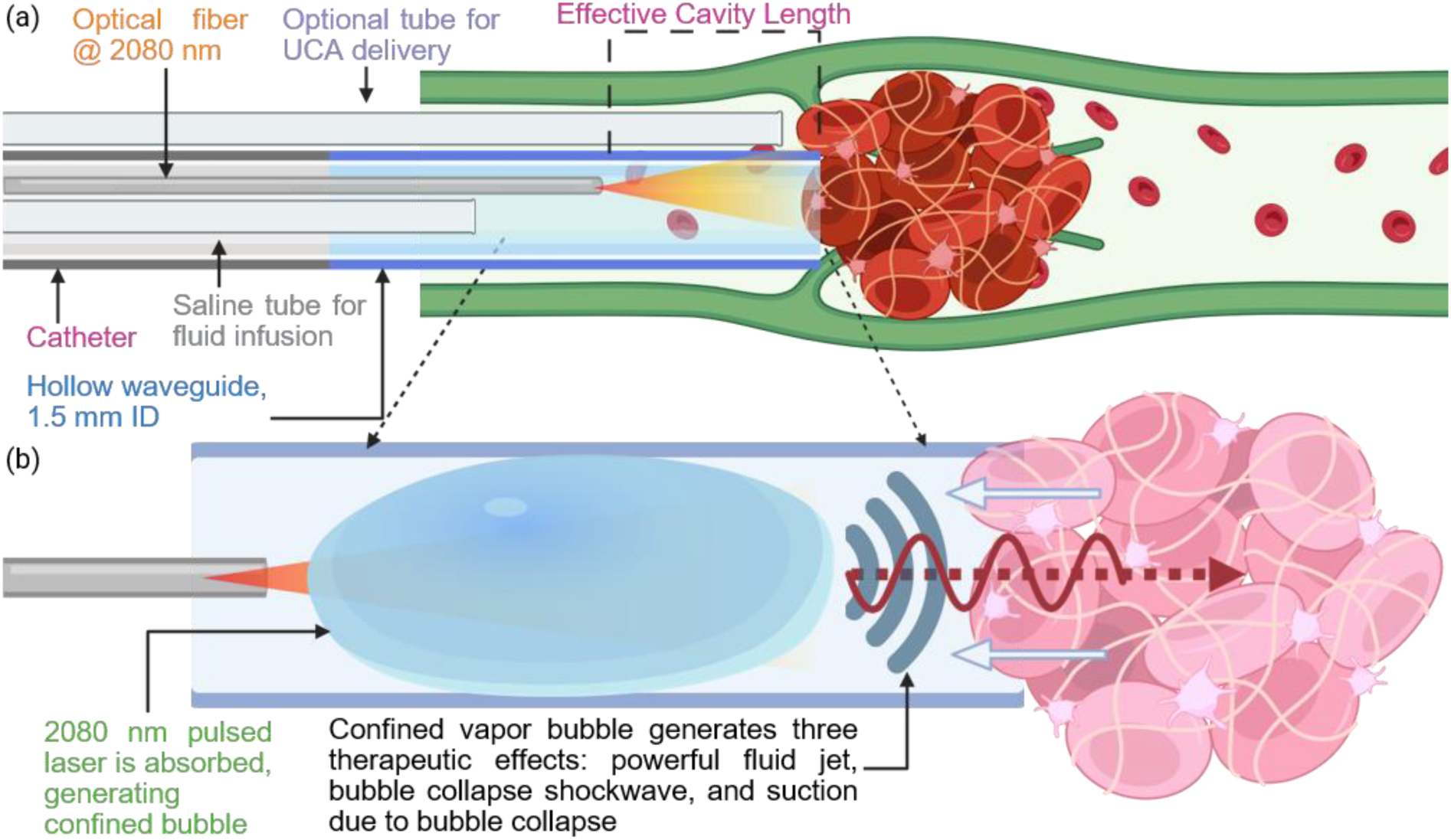
Conceptual diagram of a CLIC catheter. **a)** CLIC catheter positioned on surface of a thrombus inside an occluded vein. The catheter contains one optical fiber, delivering 2080 nm light, and a saline tube for infusion. A stainless-steel waveguide with an inner diameter (ID) of 1.5 mm is placed at the end of the catheter. An optional saline tube can be placed outside of the catheter for microbubble delivery. An effective cavity length (ECL), the distance between the optical fiber tip and waveguide end, is fixed to ensure consistent treatment efficacy. **b)** CLIC treatment mechanisms. 2080 nm light is delivered into the waveguide, vaporizing the saline inside. The vaporized saline creates a confined bubble. The formation/expansion of the bubble generates a fluid jet, while the subsequent bubble collapse generates a powerful shockwave, both of which disrupt the structure of the thrombus. The collapse also generates a low-pressure void which induces a suction effect, displacing the blood from the weakened thrombus.

Overall, CLIC shares similar principle with the research by Ohki et al. for use during neurosurgery [25], which explored the optimal ECL selection. However, differences in parameters between CLIC and Ohki’s setup yield distinct treatment mechanisms. In CLIC, treatment is initiated by delivering a 2080 nm laser pulse through the optical fiber into the saline-filled waveguide. Similar to conventional laser lithotripsy, absorption of the laser energy by the fluid leads to rapid vaporization and formation of a vapor bubble. In contrast to conventional, free-space laser lithotripsy, the presence of the surrounding waveguide constrains bubble expansion, forcing the bubble to occupy the available cavity volume. As the bubble expands, the remaining fluid within the waveguide is expelled through the outlet as a high-velocity fluid jet that penetrates the thrombus and disrupts the fibrin network. After the laser pulse terminates, the vapor bubble collapses, forming a secondary toroidal bubble that subsequently undergoes collapse. These collapse events generate shockwaves that further compromise the structural integrity of the thrombus. Lastly, the pressure drop associated with bubble collapse produces a transient suction effect that draws displaced or weakly attached blood components from the disrupted fibrin matrix into the catheter lumen or surrounding vessel.

In this study, we investigated the operational parameters and underlying mechanisms of CLIC thrombolysis using in-vitro experiments with retracted bovine clots. CLIC treatment was applied under multiple parameter combinations to evaluate thrombolytic performance. High-speed imaging was used to characterize bubble dynamics and provide experimental validation of the proposed disruption mechanisms. Treatment efficacy was quantified by measuring clot mass before and after each experiment. CLIC demonstrated substantial thrombolytic efficacy in vitro. Under the most effective parameter combination tested, retracted thrombi exhibited an average mass loss of 262.4 mg following 40 s of treatment. For comparison, recent studies of sonothrombolysis—a clinically established treatment modality for high-risk thrombi—reported mass loss rates of approximately 10.1 mg/min for retracted thrombi [26] and 53.9 mg/min for acute thrombi [27]. Under the present conditions, CLIC therefore represents an approximately 40-fold improvement over these reported values for similarly retracted thrombi.

Despite these promising results, important questions remain regarding the safety of the CLIC technique. Evaluating the size and distribution of clot debris generated during treatment is critical, as embolic fragments may pose clinical risks [26, 27]. We thus performed a systematic investigation of the treatment parameters and their influence on debris generation for assessing the safety profile of CLIC thrombolysis. Clot debris generated during experiments was collected, filtered, and analyzed to determine particle size and distribution. Together, these experiments quantify the thrombolytic efficacy of CLIC and provide evidence supporting the proposed mechanisms, indicating its potential for treating high-risk thrombi.

## 2. Methods

### 2.1 CLIC Catheter

The in vitro CLIC thrombolysis setup is shown in Fig. 2a. A silicone tube serving as a mock catheter was embedded within a 1% (w/v) agar mold containing an 8 mm diameter cylindrical channel that acted as a mock vessel. Laser pulses were generated using a commercial holmium:yttrium– aluminum–garnet (Ho:YAG) laser lithotripter (H Solvo 35 W, Dornier MedTech, Kirkland, WA, USA). The system was operated at pulse energies of 0.2 or 0.4 J per pulse with pulse durations of 78 or 106 μs and pulse repetition frequencies of 10 or 20 Hz, corresponding to dusting and fragmenting modes, respectively. Laser energy was delivered through a 365 μm core optical fiber inserted into the catheter and coupled to a stainless-steel waveguide. Both the catheter and waveguide had an internal diameter of 1.5 mm.

**Figure 2:**
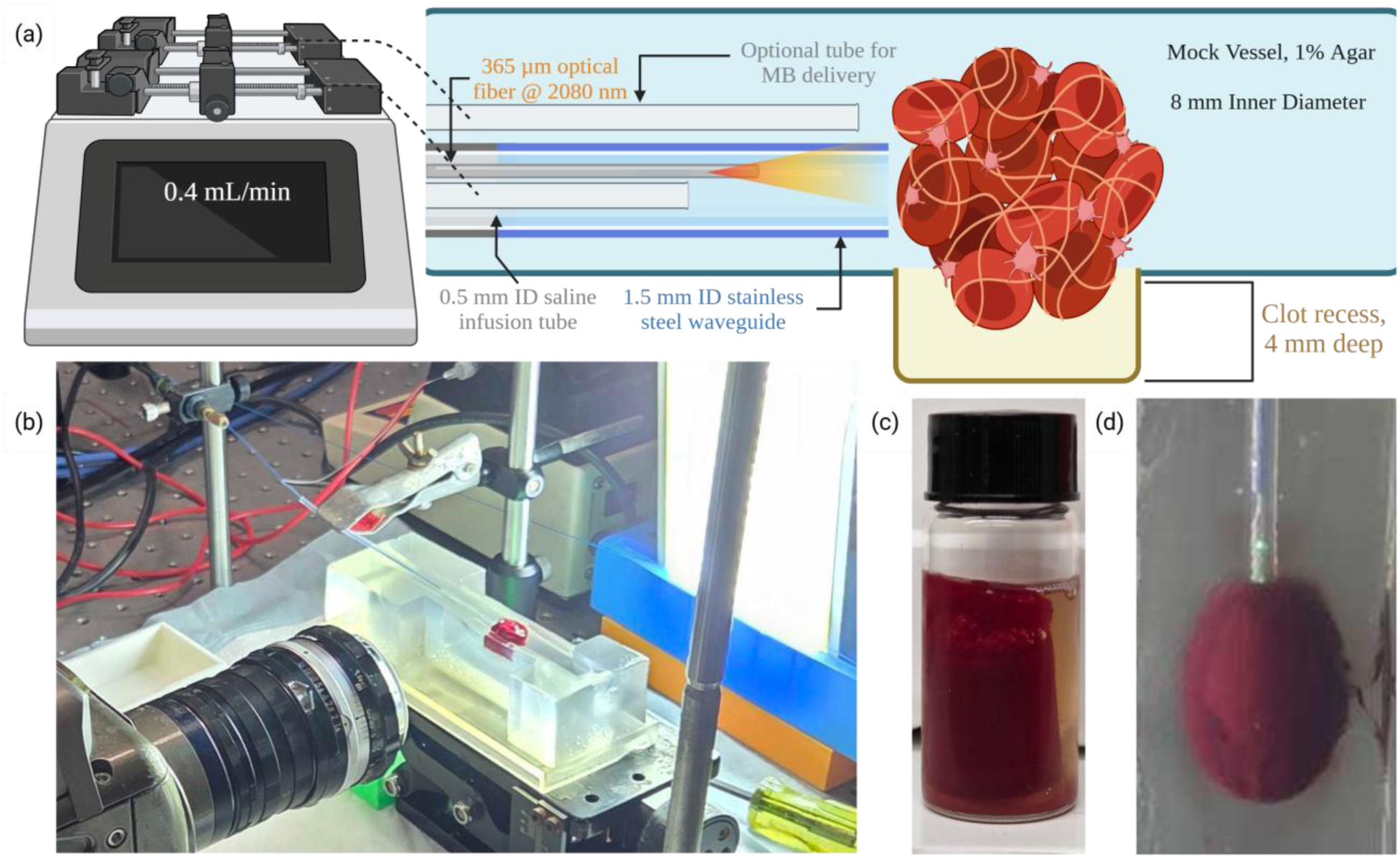
Setup for CLIC Thrombolysis Parameters Testing. **a)** Diagram of CLIC Thrombolysis testing setup. Diagram not to scale. A 1% agar mold was created with an 8 mm cylinder inside, acting as a mock vessel. A 1.5 mm ID silicone tube fitted with a 1.5 mm ID waveguide, acting as a mock catheter, was placed inside the vessel. Inside the mock catheter is a 365 µm core diameter optical fiber delivering 2080 nm light to the target, and a silicone tube (0.5 mm ID) for saline infusion. For tests including MBs, an additional silicone MB delivery tube was placed outside the catheter. Both the MB delivery tube and saline infusion tube were fed by an injection pump operating at 0.4 mL/min. ECLs of 2, 3, and 4 mm were tested for treatment efficacy. A clot recess was used to station the clot. **b)** A picture of the experimental setup with high-speed camera validation after placement of the clot. The optical fiber is fed into the mock catheter and fixed by a fiber holder. The fiber is further secured inside the mock catheter with a clamp attached to a post for stability. **c)** A bovine clot after preparation. **d)** A top-down snapshot of a bovine clot inside the experimental setup. The green pilot light seen inside of the mock catheter indicates the optical fiber tip.

In addition to the optical fiber, the catheter assembly incorporated a silicone infusion tube (305 μm ID, 635 μm OD) to deliver saline into the waveguide for laser-induced vaporization. A second silicone tube with identical dimensions was positioned alongside the catheter, with its outlet located at the catheter tip, to enable delivery of microbubbles (MBs) as required. MBs (mean diameter ≈1 μm) were produced by Vialmix agitation of a lipid solution composed of DPPC, DPPA, DPPE, and DSPE-PEG2000 (Avanti Polar Lipids) in phosphate-buffered saline containing glycerol and propylene glycol, sealed under perfluoropropane gas (C₃F₈; Fluoromed) [28]. Both infusion lines were driven by a single syringe pump operating at 0.4 mL/min. During experiments the catheter position was held fixed to isolate the effects of treatment parameters on CLIC mechanics and thrombolysis efficacy.

### 2.2 CLIC Thrombolysis Parameter Testing

Thrombolysis experiments targeted retracted bovine clots placed within the agar vessel phantom (Fig. 2b). Prior to treatment, clots were gently pressed against an absorbent pad to remove excess surface fluid and weighed. Each clot was then placed within a recessed cavity in the agar mold designed to immobilize the sample during treatment. The waveguide tip was positioned in direct contact with the clot surface, corresponding to a standoff distance (SD, here defined as the distance between the waveguide tip and target surface) of 0 mm. The ends of the mock vessel were sealed to prevent saline leakage during experiments. Apart from the saline delivered through the infusion tube, no additional fluid flow was introduced during testing to minimize any confounding variables associated with bulk flow or debris transport.

For each experiment, an ECL was selected and fixed beforehand to maintain treatment efficacy. ECL values of 2, 3, and 4 mm were selected based on preliminary high-speed imaging, which indicated that the initial laser setting (0.4 J per pulse, 106 μs pulse duration) produced bubbles approximately 2–4 mm in diameter depending on fluid temperature and experimental time point. Each clot was treated for 40 seconds, corresponding to 400 pulses in fragmenting mode or 800 pulses in dusting mode, while saline infusion was maintained. Following treatment, clots were removed, again pressed against an absorbent pad to remove excess fluid, and reweighed.

Thrombolysis efficacy was quantified as the difference between pre- and post-treatment clot mass. After each experiment, the agar mold was drained to collect dislodged clot debris for further analysis.

### 2.3 Blood Clot Preparation

The blood clots used for thrombolysis testing were prepared with a similar protocol to our previous works [20, 29]. Briefly, 8 mL of bovine blood (Lampire, Pipesville, Pennsylvania) was transferred into 10 mL borosilicate glass vials and mixed with 0.8 mL of 2.75% W/V calcium chloride solution to induce coagulation. After mixing, the blood-CaCl2 solution was placed into a 37 °C water bath to rest for 3 hours. After the clots were formed, the glass vials were placed into a 4 °C refrigerator to rest for 72 hours. The combination of the borosilicate glass vials with the refrigeration is intended to induce retraction, creating thrombi which are closer in mechanical properties to higher-risk chronic thrombi, which are typically fibrin-rich, aged, and mechanically stiffer than acute thrombi [2]. The resulting thrombi were cylindrical in shape, with approximate dimensions of 15 mm in diameter and 24 mm in length (Fig 2c) before being cut into two equal pieces. Each CLIC thrombolysis experiment tested one of these cut thrombi (Fig 2d).

### 2.4 High-Speed Photography Validation

High-speed imaging was performed to characterize bubble dynamics during CLIC operation. Imaging was conducted using a Phantom v7.3 high-speed camera (Vision Research, Wayne, NJ, USA) operating at 10,000 frames per second. Illumination was provided by a diffused LED backlight. For imaging experiments the stainless-steel waveguide was removed, and the optical fiber was retracted within the silicone catheter to reproduce different ECL values, a configuration referred to here as soft-cavity CLIC. In preliminary imaging experiments designed to observe bubble formation in the absence of a target, saline infusion was not used. Under these conditions, progressive heating of the fluid within the catheter during repeated laser pulses altered bubble size over time, as elevated temperature reduced the energy required for vaporization. In subsequent validation experiments investigating interactions between the bubble and a clot target, a syringe pump delivered room-temperature saline to stabilize the fluid temperature and maintain consistent bubble dynamics. High-speed imaging allowed visualization of early-stage bubble formation and collapse. However, middle- and late-stage CLIC mechanisms could not be observed because displaced blood and debris obscured the optical path.

### 2.5 Blood Clot Debris Analysis

Clot debris generated during treatment was collected and analyzed to evaluate particle size distribution. For venous thrombi, fragments exceeding approximately 1 mm in length are considered potentially hazardous because they may lead to pulmonary embolism [8–10]. Debris was therefore filtered and imaged on a 100 μm mesh filter. A custom MATLAB program was developed to quantify debris size from digital images. The analysis pipeline employed Otsu thresholding [30] to segment debris particles from the background, with optional watershed segmentation to separate touching or overlapping fragments and manual editing to separate any remaining erroneously connected fragments. Individual debris components were then identified and analyzed to extract geometric properties including projected area, equivalent circular diameter, and major and minor axis lengths.

### 2.6 Computational Multiphysics Analysis

Each pulse in the CLIC process involves a highly transient multiphysics interaction between laser radiation, phase transition (i.e., vaporization), multiphase fluid dynamics, and soft-tissue mechanics. While bubble dynamics can be characterized using high-speed imaging (Sec. 2.4), the transient pressure, temperature, and velocity fields within the waveguide and near the blood clot are difficult to measure experimentally. This challenge arises from the small diameter of the laser fiber, the confined geometry of the waveguide, and the large spatial and temporal gradients of these physical quantities. As a result, it is difficult to establish the causal relationship between laser radiation, bubble dynamics, and treatment outcomes through laboratory experiments alone.

In this work, physics-based numerical simulations are performed to gain insight into the physical processes occurring during each laser pulse. Specifically, the recently developed computational multiphysics framework M2C is employed to simulate laser-induced cavitation within the confined environment of the CLIC waveguide. M2C is a parallel, open-source C++ code that solves a coupled system comprised of the Navier–Stokes equations for fluid dynamics, a laser-radiation equation that models the transmission and absorption of laser energy, the latent-heat-reservoir method for predicting laser-induced cavitation, and a level-set method for tracking bubble dynamics [31, 32]. Unlike conventional bubble dynamics simulations, which assume a pre-existing bubble as the initial condition and model only its subsequent expansion and collapse, M2C resolves the full multiphysics process beginning with the onset of laser radiation. The framework has previously been applied to laser-induced cavitation problems involving pulse durations on the order of 100 μs [33] as well as nanosecond-scale pulses [34]. In both cases, the predicted bubble morphology showed close agreement with high-speed imaging measurements, while providing detailed information on laser irradiance, temperature, pressure, and velocity fields. In the latter, the simulated pressure transients also compared favorably with hydrophone measurements at probe locations [34].

The physical model employed in the present work largely follows that described in [33], including the treatment of laser fiber, laser energy absorption, and fluid thermodynamics. The simulation geometry was constructed to match the current CLIC configuration. It should be noted that the modeling of soft matter in general, and blood clots in particular, remains an active area of research. To reduce modeling complexity and isolate the fundamental cavitation physics, the simulations presented here were performed without a blood clot.

## 3. Results

### 3.1 Multiphysics Simulation Results of CLIC Thrombolysis

Fig. 3 presents the multiphysics simulation results obtained using M2C, showing the laser irradiance, temperature, pressure, and velocity fields at five representative time instants. Although CLIC involves a coupled optical–thermal–mechanical process, the simulations—performed here without a clot—clearly indicate that energy output by the CLIC catheter is primarily mechanical loading rather than heat, most notably via the fluid jet generated during bubble expansion.

**Figure 3:**
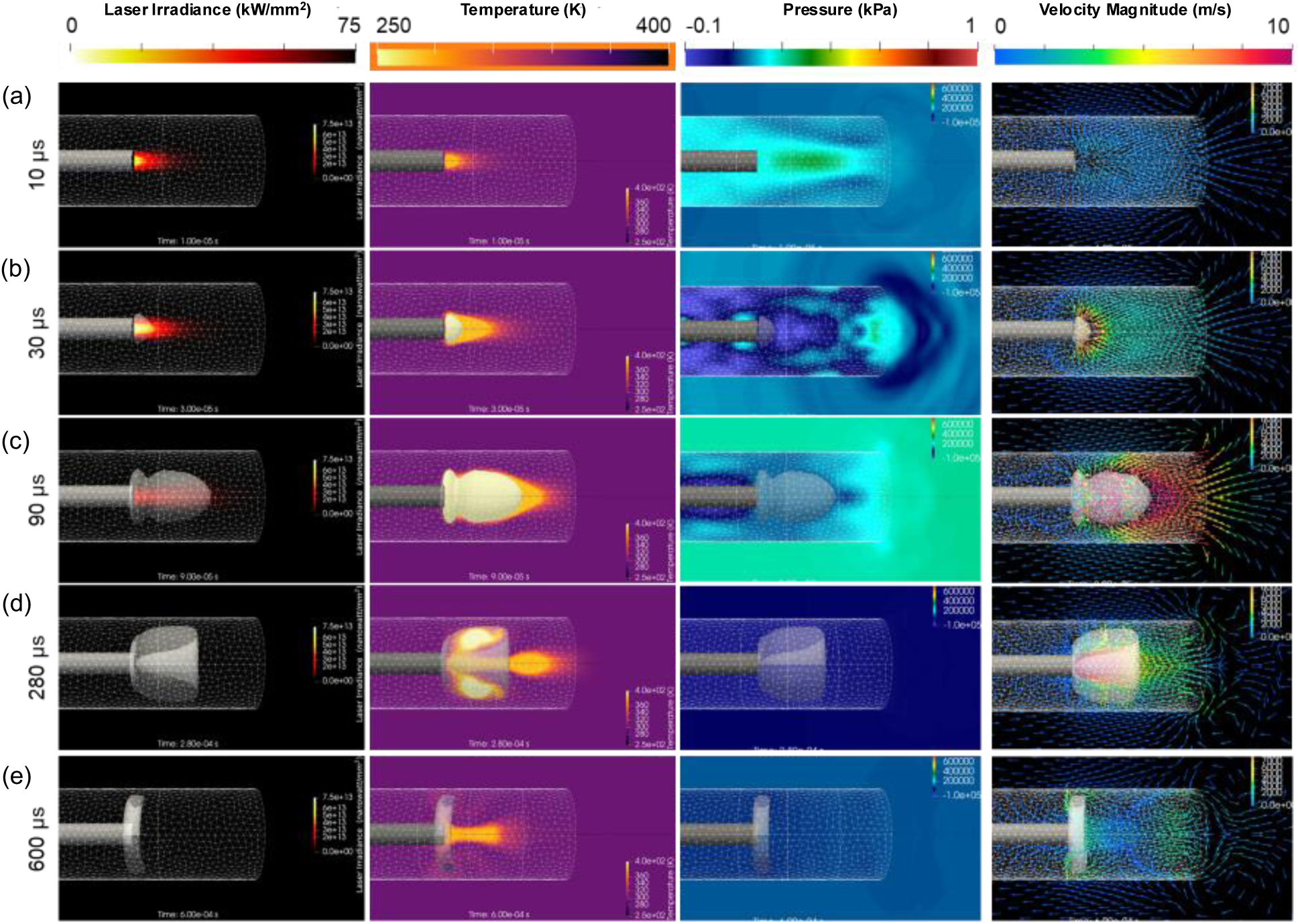
Snapshots of M2C simulation results illustrating laser-induced bubble generation, bubble growth, fluid jetting, and bubble collapse inside a mock catheter with a 2 mm ECL in the absence of a target. The catheter is represented by a triangulated surface mesh. From left to right, the laser irradiance, fluid temperature, pressure, and velocity fields are shown. Laser radiation begins at 0 μs and lasts approximately 140 μs. **a)** At 10 μs, a small volume of water in front of the laser fiber tip has reached the vaporization temperature (373 K), although cavitation has not yet occurred. **b)** By 30 μs, a small conical vapor bubble has formed within the superheated region ahead of the fiber tip. **c)** The 90 μs snapshots capture the rapid expansion of the vapor bubble, producing a high-speed jet with velocities approaching 10 m/s at the catheter outlet. **d)** At 280 μs, laser irradiation has ceased and the vapor bubble is undergoing collapse. In the absence of a target (e.g., a blood clot), the bubble collapses toward the laser fiber tip. Bubble contraction induces an inward (suction) flow at the catheter outlet with velocities of approximately 2–4 m/s. **e)** The simulation ends at 600 μs, after the bubble has collapsed into a toroidal shape.

Because the Ho:YAG laser wavelength is strongly absorbed by water, laser energy is deposited primarily near the fiber tip. Although the vapor bubble temporarily extends the optical path by allowing the laser to propagate through the vapor with negligible attenuation (also known as the “Moses effect”), the laser pulse terminates before the bubble reaches its maximum length. Therefore, laser radiation does not directly reach the catheter outlet, suggesting that the blood clot is unlikely to receive direct laser irradiation. The simulations further indicate that, although the vapor temperature inside the bubble can exceed 400 K, the liquid temperature at the catheter outlet remains below 320 K for the vast majority of the process (the initial liquid temperature is 300 K). Therefore, only limited thermal loading is expected at the clot surface.

In contrast, the mechanical effects are substantial. During bubble expansion, the fluid jet exiting the catheter reaches velocities of 5–10 m/s, approximately 5–10 times greater than the peak blood velocity in major arteries [35]. The simulations also provide detailed visualization of the jet evolution and the subsequent suction flow induced during bubble collapse. At the waveguide opening, the flow speed reaches approximately 3–6 m/s. These simulations clarify the role of the waveguide and the magnitude of the fluid jet. The simulations also visualize the suction flow examined next by high-speed imaging.

### 3.2 High-Speed Photography of 3 mm ECL CLIC Thrombolysis

High-speed photography was used to characterize bubble dynamics during CLIC. The imaging indicated that the laser-induced bubble diameter ranged from approximately 2 to 4 mm, with the observed size varying according to the temperature of the fluid within the waveguide and the applied laser energy. Subsequent validation experiments employed a controlled injection pump to maintain stable fluid temperature within the waveguide between pulses, resulting in more consistent bubble formation. A representative frame at the maximum bubble expansion is shown in Fig. 4a, where the bubble grows predominantly toward the clot surface while also expanding slightly behind the fiber tip.

**Figure 4:**
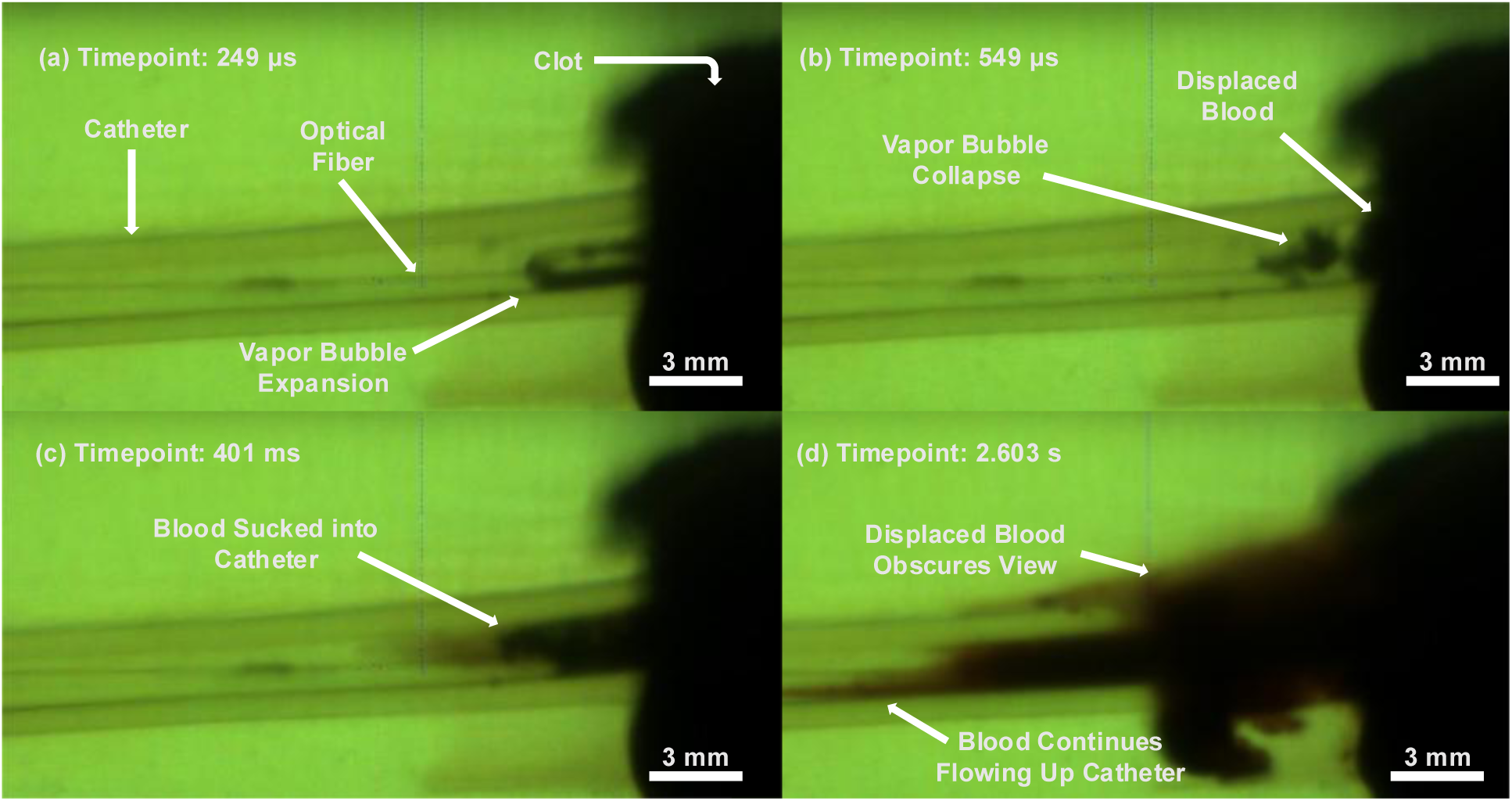
Clot therapy with high-speed photography validation. The stainless-steel waveguide was substituted by pulling the optical fiber back within a transparent silicon tube to enable high-speed camera (HSC) validation. **a)** A snapshot showing the generated bubble in a 3 mm ECL experiment. The bubble expands mostly in the direction of the clot but also expands a little towards the left, behind the fiber tip. **b)** A snapshot of the same, now collapsing bubble. The bubble collapses towards the clot surface. **c)** A snapshot of displaced blood being sucked into the catheter after several bubble collapses due to the negative pressure from the generated void. **d)** After multiple pulses, the dislodged clot debris fills the catheter and the vessel lumen.

Although the generated fluid jet could not be directly visualized in the high-speed images, it is expected to emerge from the waveguide tip and mechanically disrupt the fibrin network of the clot. Such jet-induced disruption would loosen the dense clot structure, facilitating both bubble expansion and MB infiltration into the thrombus. Following maximum expansion, the bubble collapses, producing a smaller toroidal bubble at the clot interface (Fig. 4b). In short-pulse laser lithotripsy, the collapse of this toroidal bubble—rather than the initial cavitation bubble—is understood to generate the strong shockwaves responsible for stone disruption [15, 18]. These collapse-generated shockwaves may also trigger the secondary collapse of cavitation bubbles that have infiltrated deep into the fibrin netting. Nevertheless, the high-speed camera cannot capture the bubble collapse shockwave and propagation.

Fig. 4c shows displacement of clot debris toward the catheter lumen following bubble collapse, consistent with a suction effect generated during the collapse phase. The possibility that toroidal bubble collapse may produce additional mechanical interactions with the clot, such as secondary jet formation, remains to be explored. As treatment progresses, a crater forms at the clot surface (Fig. 5). Subsequently, the increasing opening allows displaced clot debris to escape into the mock vessel lumen in addition to the catheter.

**Figure 5:**
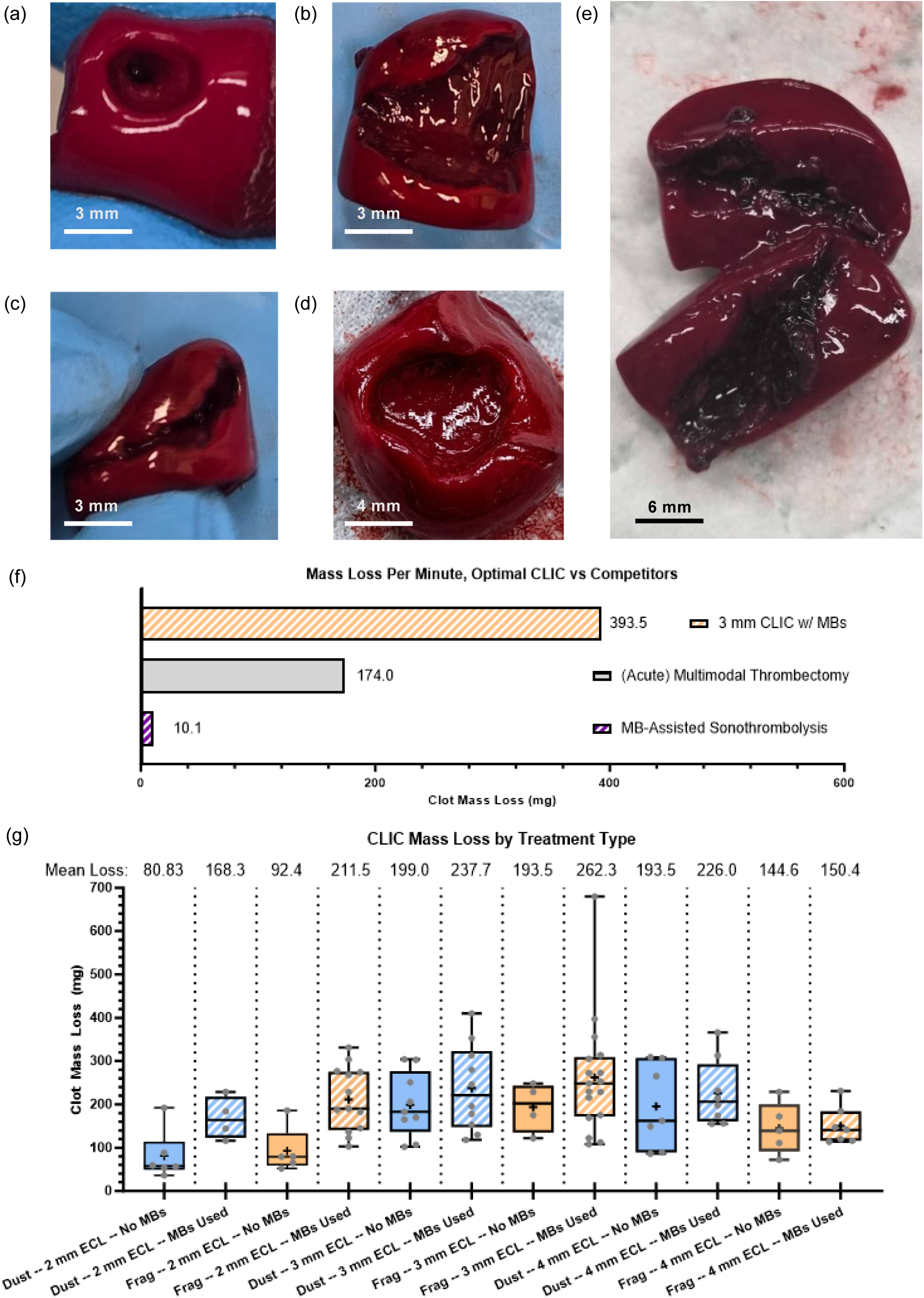
CLIC thrombolysis results demonstrating CLIC efficacy and safety for different parameter sets. **a)** Photo of clot with fragmenting (Frag) CLIC treatment with a 2 mm ECL and without MBs. The small ECL likely resulted in a smaller generated bubble, minimizing the impact from the bubble collapse shockwaves. This resulted in a shallower crater. The deeper, cylindrical hole is likely due to the initial fluid jet. **b)** Photo of an effective 4 mm fragmenting case, with a mass loss of 231 milligrams. In this case, the generated crater was deep and wide, and the clot, once removed and placed on its “back”, split apart due to the weakened clot wall. Note the complete lack of char on the clot surface, indicating that there was no thermal effect upon the clot itself. **c)** Photo of the same treated clot from 5b being restored to its original structure. **d) Photo of** a clot treated with 3 mm ECL, MB-assisted CLIC. Due to the presence of MBs during treatment, a large crater shape is generated on the surface of the clot. **e)** Photo of charred clots after 2 mm ECL CLIC. The charred regions extend deep into the clot, rather than being confined to the region around the waveguide tip. **f)** Comparisons of thrombolysis efficacy between CLIC with optimized parameters, reported MB-assisted sonothrombolysis, and reported multimodal thrombectomy. Note that the thrombectomy benchmark reflects treatment of acute clots, whereas CLIC and sonothrombolysis targeted chronic clot models. Because chronic thrombi are mechanically resistant, thrombectomy performance would be considerably reduced under comparable conditions. **g)** Box-and-whisker plots of 2, 3, and 4 mm ECL CLIC thrombolysis results. Treatment conducted with MBs present higher treatment efficacy than without MBs. Mass loss for 3 mm ECL is greater than 4 mm ECL under equivalent laser and MB conditions. For 3 mm ECL fragmenting with MBs, the average mass loss without the 680 mg outlier drops to 236.2 mg.

### 3.3 CLIC Mechanisms and Efficacy

Post-treatment thrombi provide direct visual evidence of CLIC-induced clot disruption (Fig. 5a–d). Fig. 5a shows a thrombus following a fragmentation experiment performed without MBs using a 2 mm ECL. The clot surface exhibits a shallow crater, wherein a narrower and deeper channel is visible. The shallow deformation is consistent with disruption by bubble-collapse shockwaves, whereas the deeper channel is consistent with fluid-jet penetration. The restricted crater depth may reflect the shorter ECL, which likely limited bubble expansion and consequently reduced the strength of the collapse-generated shockwave. The morphology in Fig. 5a therefore provides visual evidence of two distinct CLIC mechanisms—shockwave-driven surface disruption and jet-induced penetration—acting on the clot structure at the same time.

Fig. 5b and 5c show a thrombus treated without MBs using a 4 mm ECL. Under these conditions, the clot exhibited substantially greater disruption than typically observed for this parameter set, with a measured mass loss of 231 mg compared with a mean mass loss of 137.8 mg. Extensive removal of internal clot material from the central region of the thrombus was observed by imaging (Fig. 5b). When restored to its original shape (Fig. 5c), the external crater diameter appears comparable to that observed in Fig. 5a, although the internal disruption extends much deeper into the clot. No visible charring was observed along the interior surfaces of the disrupted thrombus. The lack of char indicates the mass loss in Figs. 5b and 5c was due to the fluid jet and bubble-collapse shockwave mechanisms. This is highly relevant to safety, as thrombus charring has been associated with increased risk of distal embolization and pulmonary embolism [36, 37]. In Fig. 5d, a large crater is observed on the clot surface. Such craters were detected exclusively during MB–assisted CLIC, supporting the hypothesis that MBs enhance the therapeutic efficacy of shockwave collapse. Clots exhibiting these large craters—observed intermittently following MB-assisted CLIC—displayed mass losses ranging from 331 to 680 mg, substantially exceeding the mean mass loss measured under any other tested condition.

Fig. 5e shows pronounced charring following 2 mm ECL CLIC. We hypothesize that the reduced ECL limits the amount of fluid within the waveguide available to absorb the incident laser energy. Consequently, excess optical energy is deposited directly into the clot, resulting in thermal ablation and surface charring. The char in Fig. 5e extends across the entire surface of the CLIC-generated tunnel within the clot, rather than being confined to the immediate vicinity of the waveguide tip. The relatively high incidence of charring during 2 mm ECL CLIC without MB assistance (approximately 80% of experiments), independent of laser mode, suggests that this configuration poses an elevated safety risk for clinical application.

### 3.4 CLIC Thrombolysis Mass Loss Results

Fig. 5f shows that CLIC demonstrated substantially higher thrombolysis rates than previously reported ultrasound- and catheter-based approaches. Under optimized conditions, CLIC achieved mass loss at a rate of 393.5 mg/min. This value exceeds the mass loss reported for MB-assisted sonothrombolysis (10.1 mg/min) [26] by nearly 40-fold and is more than twice that reported for a multimodal mechanical thrombectomy system (174 mg/min) [38]. Notably, the thrombectomy benchmark was obtained using acute thrombi, whereas both the CLIC and sonothrombolysis results correspond to chronic clot models, which are generally more mechanically resistant. Therefore, we expect that the relative performance advantage of CLIC may be even greater when compared to thrombectomy under equivalent chronic-clot conditions.

Fig. 5g shows post-treatment mass losses for different CLIC settings at 2, 3, and 4 mm ECLs. The 3 mm ECL CLIC was more effective than 2 or 4 mm ECL CLIC for the same laser mode and MB-assistance settings. We believe that the 3 mm ECL reaches an optimal balance between space available for bubble expansion (affecting the magnitude of the bubble collapse shockwaves and the velocity of the fluid jet) and ensuring that the initial bubble collapses on the target surface (which enables optimal suction/fluid dynamics).

MB assistance increased CLIC efficacy in almost all cases, the exception being 4 mm ECL fragmenting, where mean mass loss rose by only 6 mg. The difference for 3 mm ECL fragmenting is large but is driven by a single 680 mg outlier (the large crater noted above). Excluding this outlier, the 3 mm ECL MB-assisted fragmenting average falls to 236.2 mg, again comparable to dusting mode. Treatment efficacy for a 3 mm ECL is close to identical for dusting and fragmenting modes, regardless of MB usage. This is not the case at 2 or 4 mm ECL: at 2 mm ECL, fragmenting is more effective, whereas at 4 mm ECL dusting is more effective. This relationship warrants further investigation.

### 3.5 Clot Debris Analysis

The debris analysis pipeline used to quantify the number and distribution after CLIC experiments is shown in Fig. 6a-d. Briefly, an optical image of debris on top of a 100 μm mesh filter is selected for analysis (Fig. 6a). Next, an ROI is selected and segmentation parameters are chosen (Fig. 6b). A binary mask of debris particles is then extracted (Fig. 6c), which can be referenced with the original image in order to identify any erroneously outlined collection of particles for manual correction. The result is shown in Fig. 6d, with the ROI outlined in light blue and the individual particles outlined in bright yellow, allowing every particle to be tracked and measured.

**Figure 6:**
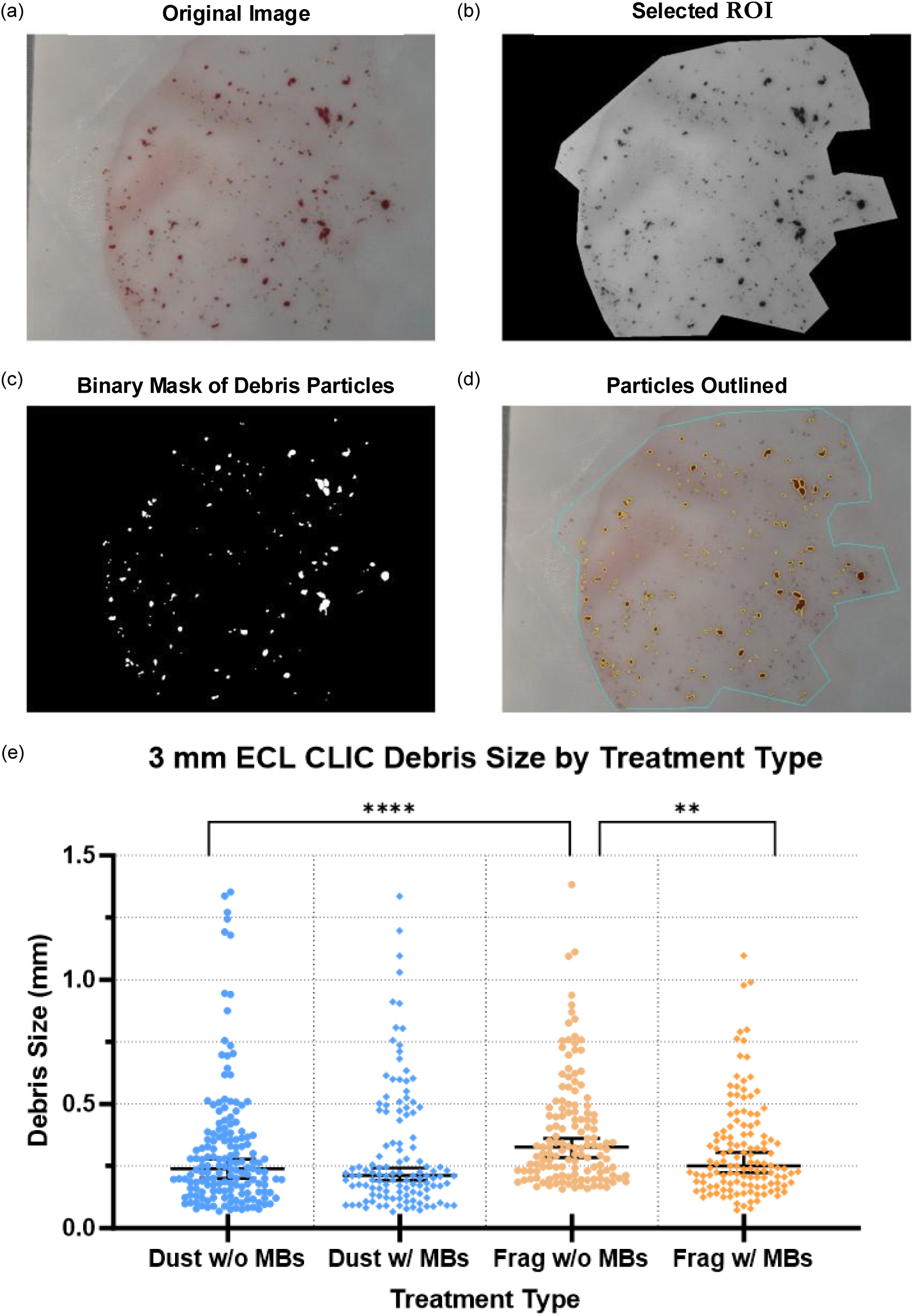
CLIC Debris Analysis. **a)** An optical image is taken of blood clot debris on top of a 100 μm filter. Care was taken to spread out the particles over the filter to ensure that blood debris did not clump together once on the filter. **b)** A region of interest is selected to analyze. **c)** A binary mask of debris particles, indicating which regions of the original image were classified as particles of sufficient area. **d)** The overlay of the segmented debris on top of the original image. Individual particles, as seen from the binary mask, are shown here outlined in thick bright yellow. **e)** Analysis of post-treatment debris from different types of 3 mm ECL. Two-tailed Mann–Whitney tests with Bonferroni correction (α = 0.0125). Frag w/o MB vs Dust w/o MB, P < 0.0001 (Hodges–Lehmann median difference, −0.082 mm); Frag w/o MB vs Frag w/ MB, P = 0.0010 (−0.059 mm); Dust w/o MB vs Dust w/ MB, P = 0.563 (0.009 mm); Frag w/ MB vs Dust w/ MB, P = 0.075 (0.03213 mm). n = 135, 159, 120 and 129 for Frag w/o MB, Dust w/o MB, Dust w/ MB and Frag w/ MB, respectively.

Debris generated during CLIC remained predominantly sub-millimeter in size, indicating a low propensity for large embolic fragment formation. Across all conditions tested at a 3 mm ECL, the maximum observed particle diameter remained below 1.4 mm (Fig. 6e). Fragmenting without MBs produced the largest debris overall (median 0.325 mm), significantly larger than debris generated by dusting without MBs (median 0.239 mm, P < 0.0001) and by fragmenting with MBs (median 0.251 mm, P = 0.001). In contrast, MB use did not significantly alter debris size in dusting mode (median 0.212 mm versus 0.239 mm, P = 0.563), and debris distributions were also not significantly different between dusting and fragmenting when MBs were present (P = 0.075).

## 4. Discussion

### 4.1 CLIC Treatment Mechanisms and Safety

In this paper, we have demonstrated that constrained laser-induced cavitation is an effective catheter-based modality for treating deep vein thrombosis. We demonstrated CLIC achieved 40-fold improvement in mass-loss rate over competing modalities such as sonothrombolysis [26] under comparable conditions and doubled the mass-loss of a multimodal thrombectomy system treating an acute clot [38]. We have explored the mechanisms of CLIC thrombolysis, namely the fluid jet disruption of the clot fibrin netting, the bubble collapse, resulting in strong shockwaves into the clot and further disrupting its internal structure, and the suction effect from the low-pressure void induced by the bubble collapse within the waveguide. We collected supporting evidence through simulations and high-speed imaging. However, questions remain regarding the mechanisms of middle- and late-stage CLIC thrombolysis, especially regarding a) how the expanding space inside the clot in front of the waveguide—i.e., the crater—affects bubble size and subsequently increases shockwave potency, and b) the role of the fluid jet after the initial cutting of the fibrin structure, especially regarding the displacement of now-loose blood cells and small clot debris.

The ECL strongly influences treatment efficacy. Guided by our mechanistic understanding and the literature [25, 39, 40], the ECL primarily governs whether the laser-generated vapor bubble can extend beyond the waveguide, maintain contact with the clot surface, and collapse in a geometry favorable for effective clot disruption. At an intermediate ECL of 3 mm, these conditions are likely to be best balanced: the bubble can spill out of the waveguide [26], which ensures that the bubble will collapse towards the clot surface and form a secondary toroidal bubble which, as mentioned in section 3.2, generates a therapeutic shockwave effect. Although toroidal bubble collapse is a dominant therapeutic mechanism in laser lithotripsy, its role in CLIC thrombolysis remains uncertain. First, the thrombi are weaker and more elastic than the kidney stones treated by laser lithotripsy. Second, the post-treatment thrombi consistently show a cylindrical tunnel, regardless of MB use, which is most readily explained by the generated fluid jet. Third, the waveguide should make the maximum initial bubble in CLIC much smaller than in free-space laser lithotripsy, which in turn should limit the size and potency of the toroidal bubble and subsequent collapse shockwaves.

The distinct treatment responses observed at 2 mm and 4 mm ECL indicate that CLIC efficacy depends not only on nominal bubble size, but also on how cavitation dynamics couple to the clot at different ECLs. At a 2 mm ECL, fragmenting appeared to outperform dusting, suggesting that limited space available for bubble expansion may preferentially limit the effectiveness of lower-energy, higher repetition-rate dusting mode pulses. In contrast, at 4 mm ECL, dusting appeared to perform better than fragmenting, suggesting that the dominant treatment processes may shift as the ECL increases. One possible explanation is that once the bubble no longer consistently extends to the clot surface, clot-directed collapse effects become less prominent and treatment outcome depends more strongly on cumulative exposure, fluid-mediated erosion and debris clearance.

Observations from laser lithotripsy studies [15, 18] and high-speed imaging support this theory, indicating that generated vapor bubbles that fail to contact the target are expected to collapse preferentially towards the fiber rather than the target, which would further reduce any contribution from toroidal bubble-collapse shockwaves. In this regime, dusting may therefore benefit from its higher pulse repetition rate, and fragmenting may be disadvantaged if its bubble geometry is less effective at maintaining clot contact over the longer stand-off distance.

A shorter ECL (e.g., 2 mm) may also increase the risk of direct optical and thermal injury to the thrombus. When the fiber tip is positioned too close to the clot surface, a greater fraction of the laser energy may be deposited into the thrombus rather than absorbed within the saline, increasing the likelihood of local charring. This risk is further amplified if fiber positioning is unstable or the waveguide is absent. Consistent with this interpretation, clot charring was observed under short-distance conditions, indicating that sufficiently large ECLs may be required not only for efficient cavitation-mediated thrombolysis but also to minimize unintended thermal deposition and preserve procedural safety.

To date, there is no definitive size-based risk threshold for pulmonary embolism, but previous studies have suggested that pulmonary embolic risk increases considerably when generated debris size exceeds 1 mm [8–10]. Debris analysis suggests that CLIC primarily disrupts thrombus through progressive erosion rather than the detachment of large fragments. Across all treatment conditions, particle sizes remained predominantly sub-millimeter, with even the largest fragments remaining below 1.4 mm. Although in vitro measurements cannot fully reproduce in vivo transport and filtration dynamics, this size regime is generally unlikely to produce clinically significant obstruction in the venous circulation, where vessel diameters are substantially larger.

MB usage further reduced debris size in fragmenting mode, suggesting that cavitation nucleation may influence fragmentation behavior. One possible explanation is that MBs promote more distributed cavitation activity, favoring repeated small-scale erosive events. In contrast, fragmenting without MBs produced the largest debris, consistent with a regime in which higher-energy cavitation events may detach larger fragments prior to subsequent erosion. Together, these observations support a CLIC thrombolysis mechanism dominated by cavitation-driven erosion, fluid jetting and suction-mediated evacuation.

### 4.2 Improving CLIC via Movement and Guidance

A key question is how much catheter translation during treatment, particularly when paired with imaging guidance, can further increase efficacy. CLIC already greatly exceeds the efficacy of conventional methods. In the present experiments the catheter was held static rather than translated to maintain an optimal SD. Translation is therefore expected to further increase efficacy. We expect that although the growing crater will allow a larger laser-induced bubble, and thus a stronger collapse shockwave, this gain will eventually be offset by the increased SD, which will diminish treatment efficacy and eventually lead to the laser-induced bubble collapse towards the fiber rather than the clot. Moving the catheter and maintaining an optimal SD over the course of treatment should remove this issue. Although higher efficacy is not strictly required, this headroom could instead be used to lower laser energy and MB dosage—improving safety—while retaining an order-of-magnitude margin over conventional modalities. This gain should be most pronounced for acute thrombi, which are less stiff than the retracted clots studied here. We plan to evaluate catheter translation in future work.

Guiding catheter translation will require intraoperative, real-time imaging. Conventional imaging modalities are each limited in this role. CT venography would be able to locate the venous thrombi but may not provide sufficient information regarding the treatment crater growth, and furthermore uses ionizing radiation, making it challenging for continuous intraoperative imaging guidance [41, 42]. Ultrasound, while safe, capable of continuous real-time imaging guidance, will struggle with identifying a deep thrombus with sufficient contrast and may not be able to visualize the crater growth [43, 44]. Complementary Doppler ultrasound, however, can quantify real-time changes in vessel occlusion [45]. Photoacoustic imaging (PAI) is a hybrid imaging modality which combines the deep penetration and resolution of ultrasound with the molecular sensitivity of optical imaging [46–50] and is promising for intraoperative CLIC guidance.

From a practical standpoint, the CLIC catheter can accommodate a second optical fiber for imaging wavelengths, simplifying integration of PAI. The use of this imaging fiber within the catheter to directly deliver light to an internal target is a technique that has been termed “internal-illumination photoacoustic tomography”, or II-PAT [20, 51, 52]. The use of an ultrasound array sitting outside of the target system—whether an arm or leg, in an in-vivo setting, or a mock vessel, as in our current in-vitro setup—would allow for capture of the II-PAT signals and also allow for the previously mentioned Doppler ultrasound, which would allow for co-registered monitoring of blood flow changes during treatment. Unlike ultrasound, II-PAT can localize the thrombus, as we have demonstrated previously [20, 29]. Additionally, because the light will be directly delivered to the target as opposed to traveling through centimeters of tissue, the pulse energy required for a strong photoacoustic signal should be low, reducing the risk of thermal damage. Furthermore, II-PAT is uniquely capable of providing deep molecular information through the use of NIR-II (1000-1700 nm) wavelengths [49, 53, 54], which allow for PAI to investigate the typical thrombus components such as lipids, collagen, and fibrin. Traditionally, due to the high optical absorption of blood and water, it is difficult to derive deep information regarding these components, especially past 1 cm depths [50, 53]. By delivering light directly to the target, II-PAT could capture clot-composition information in real time, providing more informative intraoperative guidance.

### 4.3 Outlook on CLIC

The fluid mechanics of middle- and late-stage CLIC remain to be explored, and the influence of intra-crater fluid dynamics on efficacy is currently unclear. There are several approaches we can use to observe CLIC fluid mechanics, namely particle image velocimetry (PIV) and the use of a transparent blood-mimicking phantom for use in high-speed photography. As previously mentioned, high-speed photography cannot provide information of middle- and late-stage CLIC mechanics because the presence of blood. We intend to create blood-mimicking agarose gel-based thrombus phantoms, which should be transparent and have similar physical properties to our retracted bovine thrombi [55]. PIV has been used to visualize fluid dynamics in cavitation [56, 57], and we expect that PIV will allow for visualization of the fluid dynamics through our transparent thrombus phantoms. We aim to relate this improved mechanistic understanding to the debris size and size distribution measured for each parameter set.

Finally, CLIC thrombolysis may extend to other settings, such as kidney and bladder thrombosis. Unlike DVT, kidney and bladder thrombolysis do not have strong safety risks tied to debris size, as any debris generated during treatment should be flushed out of the system rather than propagating further into the bloodstream as in arterial or venous thrombolysis [58–60]. Furthermore, the available space for catheter insertion will be much larger than most DVT cases. For these applications, CLIC may achieve faster and more effective thrombus removal than existing methods. Kidney thrombosis is a particularly viable target, since a laser lithotripter is already used for kidney-stone treatment and could be repurposed for CLIC.

## 5. Conclusion

In summary, we show that constrained laser-induced cavitation enables rapid and efficient thrombolysis of mechanically resilient clot models using a miniaturized optical platform. By confining laser-induced bubble dynamics within a waveguide, CLIC achieves substantially greater clot removal than conventional sonothrombolysis benchmarks while producing thrombus debris predominantly below a 1 mm venous embolic risk threshold. Our results further indicate that treatment performance is highly sensitive to ECL, pulse mode and MB use, underscoring the importance of cavitation–clot coupling and treatment geometry in determining therapeutic outcome. Under optimized conditions, CLIC removed an average of 262.4 mg of retracted clot in 40 seconds, almost 40-fold that of reported sonothrombolysis benchmarks under comparable in-vitro conditions. Although the relative contributions of shockwaves, fluid jetting and suction remain to be resolved more fully, these findings establish CLIC as a promising framework for high-efficacy, catheter-based thrombolysis. With further optimization of imaging guidance, catheter translation, and safety validation, CLIC may expand the treatment of resistant venous thrombi while enabling less invasive intervention in anatomically challenging situations.

## Author Contributions

**Joseph Yang:** Conceptualization, Methodology, Investigation, Data curation, Formal analysis, Visualization, Writing – original draft, Writing – review and editing. **Daiwei Li:** Methodology, Investigation, Writing – review and editing. **Kevin Wang:** Methodology, Software, Formal analysis, Visualization, Writing – review and editing. **Pei Zhong:** Conceptualization, Methodology, Writing – review and editing. **Junjie Yao:** Conceptualization, Supervision, Project administration, Resources, Funding acquisition, Writing – review and editing.

## Funding

J.Y. thanks the support by the United States National Institutes of Health (NIH) grants RF1 NS115581, R01 NS111039, R01 EB028143, R01 DK139109, R01 DK052985, R01 MH135932, R01 ES036951; The United States National Science Foundation (NSF) CAREER award 2144788; Duke University Pratt Beyond the Horizon Grant; Eli Lilly Research Award Program; Chan Zuckerberg Initiative Grant (2020-226178 and 2024-349531); Duke University DST Spark Seed Grant; Duke Coulter Translational Grant; North Carolina Biotechnology Center Triangle Research Grant (2024-TRG-0041); American Heart Association Collaborative Science Award (25CSA1417550).

## Declarations

J.Yao has a financial interest in Lumius Imaging, Inc., and Merge Labs, which did not support this work. The other authors declare no competing interests.

